# Related enteric viruses have different requirements for host microbiota in mice

**DOI:** 10.1101/733766

**Authors:** Christopher M. Robinson, Mikal A. Woods Acevedo, Broc T. McCune, Julie K. Pfeiffer

## Abstract

Accumulating evidence suggests that intestinal bacteria promote enteric virus infection in mice. For example, previous work demonstrated that antibiotic treatment of mice prior to oral infection with poliovirus reduced viral replication and pathogenesis. Here we examined the effect of antibiotic treatment on infection with coxsackievirus B3 (CVB3), a picornavirus closely related to poliovirus. We treated mice with a mixture of five antibiotics to deplete host microbiota and examined CVB3 replication and pathogenesis following oral inoculation. We found that, like poliovirus, CVB3 shedding and pathogenesis were reduced in antibiotic-treated mice. While treatment with just two antibiotics, vancomycin and ampicillin, was sufficient to reduce CVB3 replication and pathogenesis, this treatment had no effect on poliovirus. Quantity and composition of bacterial communities were altered by treatment with the five antibiotic cocktail and by treatment with vancomycin and ampicillin. To determine whether more subtle changes in bacterial populations impact viral replication, we examined viral infection in mice treated with milder antibiotic regimens. Mice treated with one-tenth the concentration of the normal antibiotic cocktail supported replication of poliovirus but not CVB3. Importantly, a single dose of one antibiotic, streptomycin, was sufficient to reduce CVB3 shedding and pathogenesis, while having no effect on poliovirus shedding and pathogenesis. Overall, replication and pathogenesis of CVB3 is more sensitive to antibiotic treatment than poliovirus, indicating that closely related viruses may differ in their reliance on microbiota.

**Importance:** Recent data indicate that intestinal bacteria promote intestinal infection of several enteric viruses. Here we show that coxsackievirus, an enteric virus in the picornavirus family, also relies on microbiota for intestinal replication and pathogenesis. Relatively minor depletion of the microbiota was sufficient to decrease coxsackievirus infection, while poliovirus infection was unaffected. Surprisingly, a single dose of one antibiotic was sufficient to reduce coxsackievirus infection. Therefore, these data indicate that microbiota can influence enteric virus infection through distinct mechanisms, even for closely related viruses.

## Introduction

Enteric viruses are spread through the fecal-oral route and cause morbidity and mortality in humans worldwide (1–4). These viruses initiate infection in the gastrointestinal tract, which is home to a community of bacteria, fungi, and viruses, termed the microbiota. Microbiota play important roles in human development, metabolism, and immunity (5). We and others have shown that intestinal bacteria can enhance viral replication and pathogenesis of enteric viruses, including poliovirus, reovirus, rotavirus, mouse mammary tumor virus, and noroviruses (6–11). While the mechanisms underlying bacterial promotion of enteric virus replication are not entirely clear, several groups have shown that bacteria can enhance viral infection through effects on host immunity and/or effects on viral particles (12). For example, we have shown that bacterial surface polysaccharides bind to poliovirus, a member of the *Picornaviridae*, and enhance virion stability and cell attachment (13). However, it is not known whether bacteria influence the replication of other picornaviruses *in vivo* and whether mechanisms of bacterial enhancement of viral infection are conserved for multiple viruses within a viral family.

Coxsackievirus is an enteric virus in the *Picornaviridae* family that initiates infection in the gastrointestinal tract. Among enteric viruses, coxsackieviruses are very common and can cause hemorrhagic conjunctivitis, hand, foot, and mouth disease, and myocarditis (14–17). Coxsackievirus B3 (CVB3) shares 62.3% sequence identity with poliovirus, a closely related picornavirus. CVB3 is the most common virus associated with viral myocarditis (18, 19), which can lead to heart disease and heart failure. Unfortunately, there are no vaccines or treatments for coxsackievirus infections.

Previously, we established an oral inoculation model for CVB3 to study viral replication within the gastrointestinal tract (20, 21), and here we examined whether bacteria influence coxsackievirus infection. Similar to our observations with poliovirus (9), we found that intestinal bacteria enhance CVB3 fecal shedding and pathogenesis. However, CVB3 and poliovirus differed in their reliance on microbiota, since minimal depletion of the microbiota reduced replication and pathogenesis of CVB3 but not poliovirus. These data suggest that CVB3 is more sensitive than poliovirus to perturbations in the microbiota. Overall, these data indicate that closely related enteric viruses may rely on the microbiota through distinct mechanisms.

## Results

### Intestinal bacteria enhance CVB3 replication and pathogenesis *in vivo*

Previously we determined that intestinal bacteria enhance poliovirus replication and pathogenesis in orally-inoculated mice (9). To determine if bacteria enhance infection of another closely related enteric virus, we examined CVB3 replication and pathogenesis in antibiotic-treated mice. To facilitate direct comparison of poliovirus and CVB3, we performed these experiments in mice that are orally susceptible to both viruses, due to expression of the human poliovirus receptor (PVR) and lack of the interferon alpha/beta receptor (IFNAR) (C57BL/6 PVR^+/+^ IFNAR^−/−^ mice). Following weeklong administration of a combination of ampicillin, neomycin, metronidazole, vancomycin, and streptomycin, mice were perorally inoculated with of 5×10^7^ PFU of CVB3. Feces were collected at 24, 48, and 72 hours post-inoculation and CVB3 titers were determined by plaque assay. Similar to poliovirus (9), we found that mice treated with antibiotics had reduced CVB3 shedding at 48 and 72 hours post-inoculation (Fig. 1A), reduced CVB3 titers in tissues at 72 hours post-inoculation (Fig. 1B), and reduced lethality (Fig. 1C) when compared to untreated mice. These data suggest that intestinal bacteria enhance CVB3 replication and pathogenesis.

**Figure 1.**
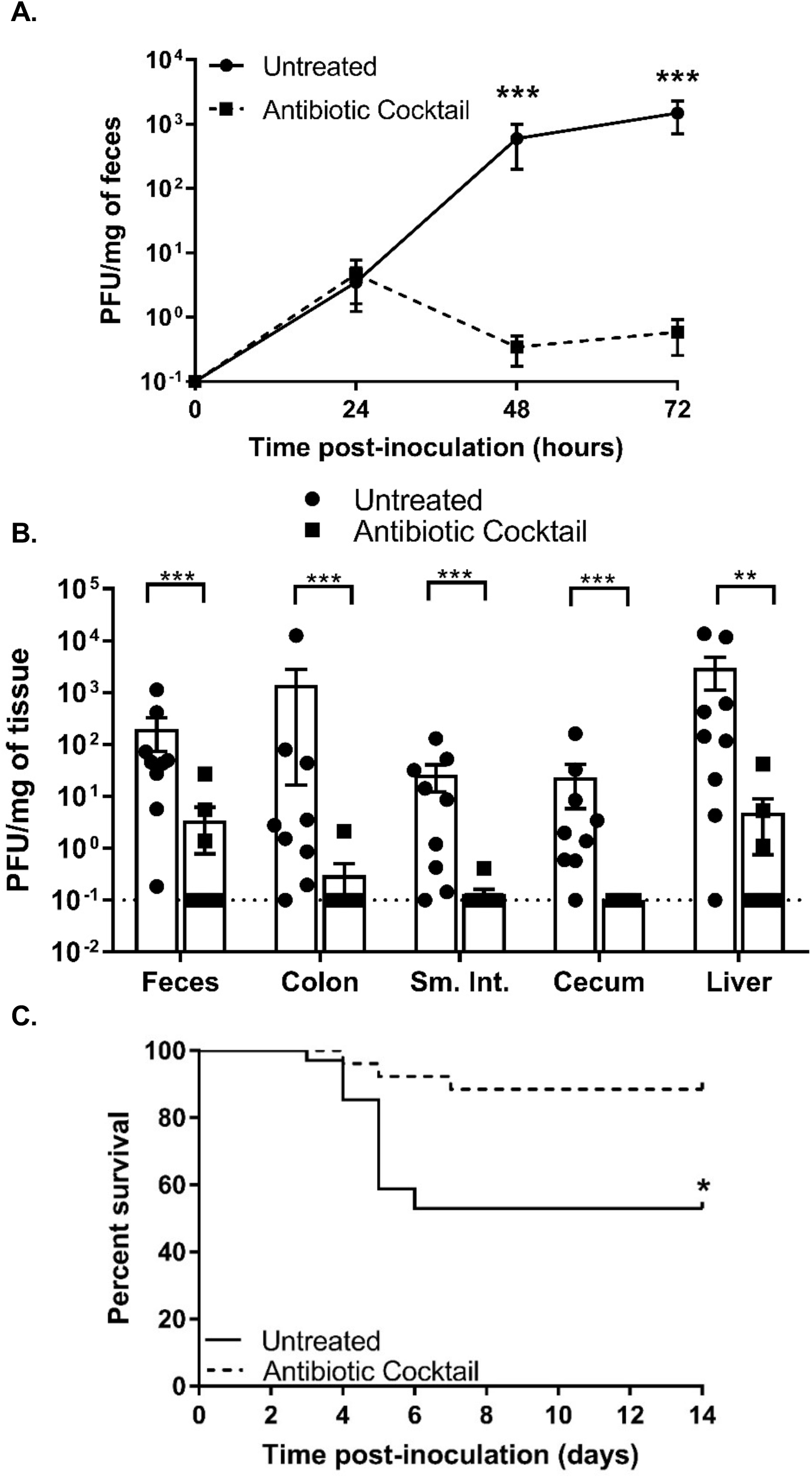
Treatment with a cocktail of five antibiotics reduces CVB3 shedding and lethality. Male C57BL/6 PVR^+/+^ IFNAR^−/−^ mice were treated with or without a combination of 5 antibiotics (ampicillin, neomycin, streptomycin, metronidazole, and vancomycin) for 5 days by oral gavage followed by *ad libitum* administration in drinking water for 9 days prior to oral inoculation with 5×10^7^ PFU of CVB3. Viral titers in feces collected at 24, 48, and 72 hours post-inoculation (A) or tissues collected at 72 hours post-inoculation (B) were determined by plaque assay. Data are mean ± SEM. *p<0.05, **p<0.001, ***p<0.0001 Mann-Whitney Test. (C) Survival of untreated or antibiotic-treated mice orally inoculated with CVB3. *p<0.05, Log-rank test. n=9-25 mice per group.

### Treatment with ampicillin and vancomycin reduces CVB3, but not poliovirus, infection

To determine the effect of specific antibiotics on CVB3 replication, we treated mice with either individual antibiotics or in different combinations prior to oral inoculation with 5×10^7^ PFU of CVB3. Feces were collected at 72 hours post-inoculation, and CVB3 titers were quantified by plaque assay. We found no difference in CVB3 fecal titers of untreated mice compared to those treated with vancomycin (Fig. 2A). In contrast, CVB3 fecal titers were reduced in mice treated with ampicillin or ampicillin plus vancomycin compared to untreated mice (Fig. 2A). In fact, fecal titers in mice treated with ampicillin and vancomycin were similar to fecal titers in mice treated with the five antibiotic cocktail. Additionally, we observed that mice treated with ampicillin and vancomycin prior to oral inoculation were protected from CVB3-induced lethality (Fig 2B). Overall, these data suggest that treatment with ampicillin and vancomycin is sufficient to reduce both CVB3 shedding and pathogenesis.

**Figure 2.**
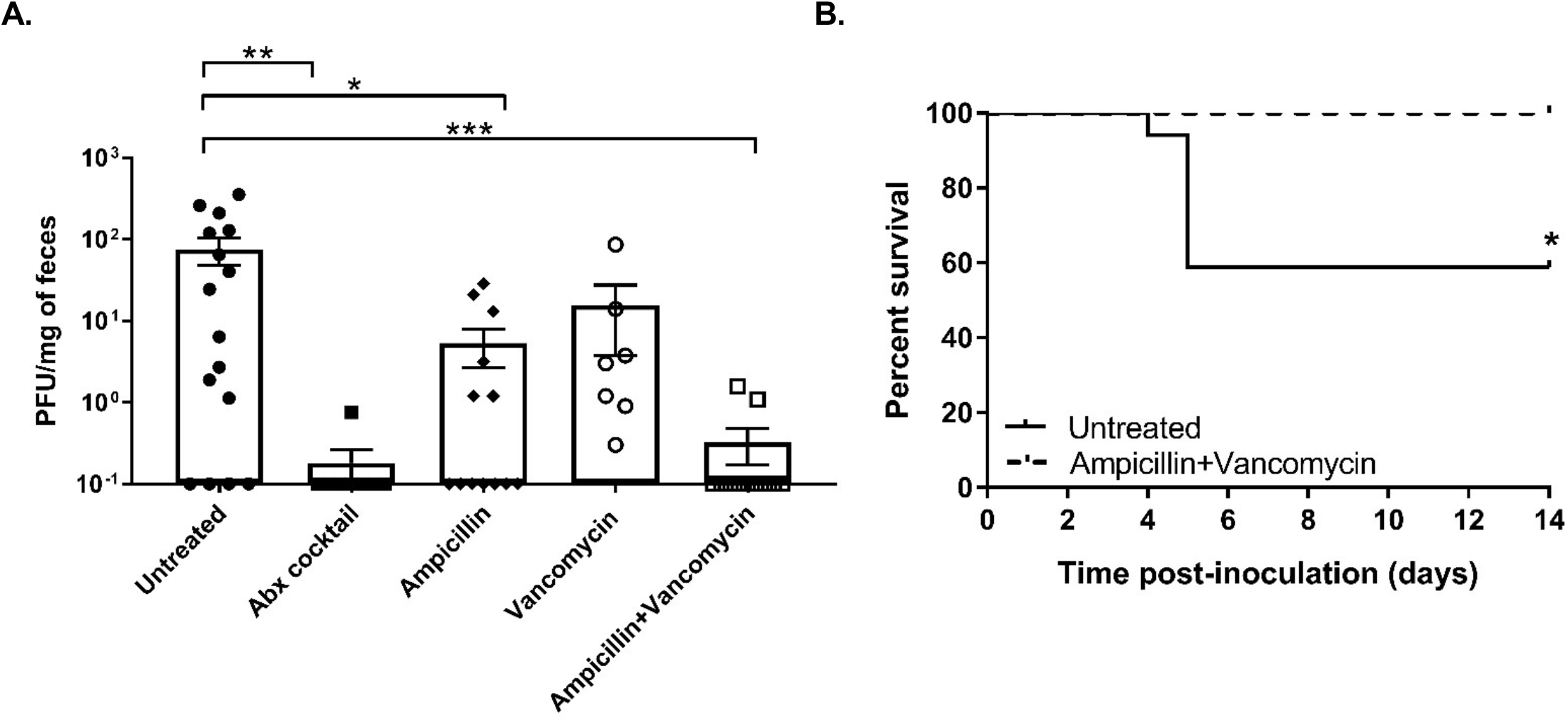
Treatment with two antibiotics, ampicillin and vancomycin, is sufficient to reduce CVB3 shedding and lethality. Male C57BL/6 PVR^+/+^ IFNAR^−/−^ mice were treated with or without various antibiotics for 5 days by oral gavage followed by *ad libitum* administration in drinking water for 9 days prior to oral inoculation with 5×10^7^ PFU of CVB3. (A) Viral titers in feces collected at 72 hours post-inoculation were determined by plaque assay. Data are mean ± SEM. *p<0.05, **p<0.001, ***p<0.0001 Mann-Whitney Test. (B) Survival of untreated or ampicillin and vancomycin-treated mice. *p<0.05, Log-rank test. n=12-17 mice per group.

Next, we examined whether treatment with ampicillin and vancomycin would be sufficient to reduce poliovirus replication and pathogenesis. Since our laboratory previously determined that depletion of intestinal bacteria with a four antibiotic cocktail reduced poliovirus replication and pathogenesis (9) and CVB3 and poliovirus are from the same viral family, we hypothesized that ampicillin and vancomycin treatment would be sufficient to reduce poliovirus infection. To address this hypothesis, we treated mice with ampicillin and vancomycin prior to oral inoculation with 2×10^7^ PFU of poliovirus. Feces were collected at 24, 48, and 72 hours post-inoculation and poliovirus titers were quantified by plaque assay. We found no significant difference in poliovirus fecal titers of untreated mice compared to those treated with ampicillin and vancomycin (Fig. 3A). These data suggest that treatment with ampicillin and vancomycin does not alter poliovirus replication in the intestine. However, high fecal titers are not always indicative of poliovirus replication in the intestine (9, 20). Input inoculum virus can be shed for a few days following infection and confound interpretation of viral titer data, particularly during antibiotic treatment due to reduced peristalsis (9, 20). Therefore, to ensure that the fecal titers detected in poliovirus-infected mice are due to replicated rather than input virus, we differentiated between inoculum and replicated viruses in mouse feces using light-sensitive viruses. Poliovirus was propagated in the presence of neutral red dye to create a light-sensitive poliovirus stock. Neutral red dye within virions confers light-sensitivity due to RNA cross-linking. Upon viral replication in the dark, such as in the gastrointestinal tract, progeny virions are light insensitive, which facilitates the assessment of replication by differentiating inoculum virus from replicated virus (9, 13, 21). Following inoculation with light-sensitive, neutral red poliovirus, feces were collected at 24, 48, and 72 hours post-infection in the dark. Following processing, poliovirus replication was quantified by determining the ratio of light-verses dark-exposed fecal titers. Again, in contrast to CVB3, we found that poliovirus replication was not reduced in mice treated with ampicillin and vancomycin (Fig. 3B). Furthermore, treatment of mice with ampicillin and vancomycin did not alter the survival of mice inoculated with poliovirus (Fig. 3C). Overall these data suggest that treatment with ampicillin and vancomycin is sufficient to reduce CVB3 shedding and pathogenesis, while having no effect on poliovirus replication and pathogenesis.

**Figure 3.**
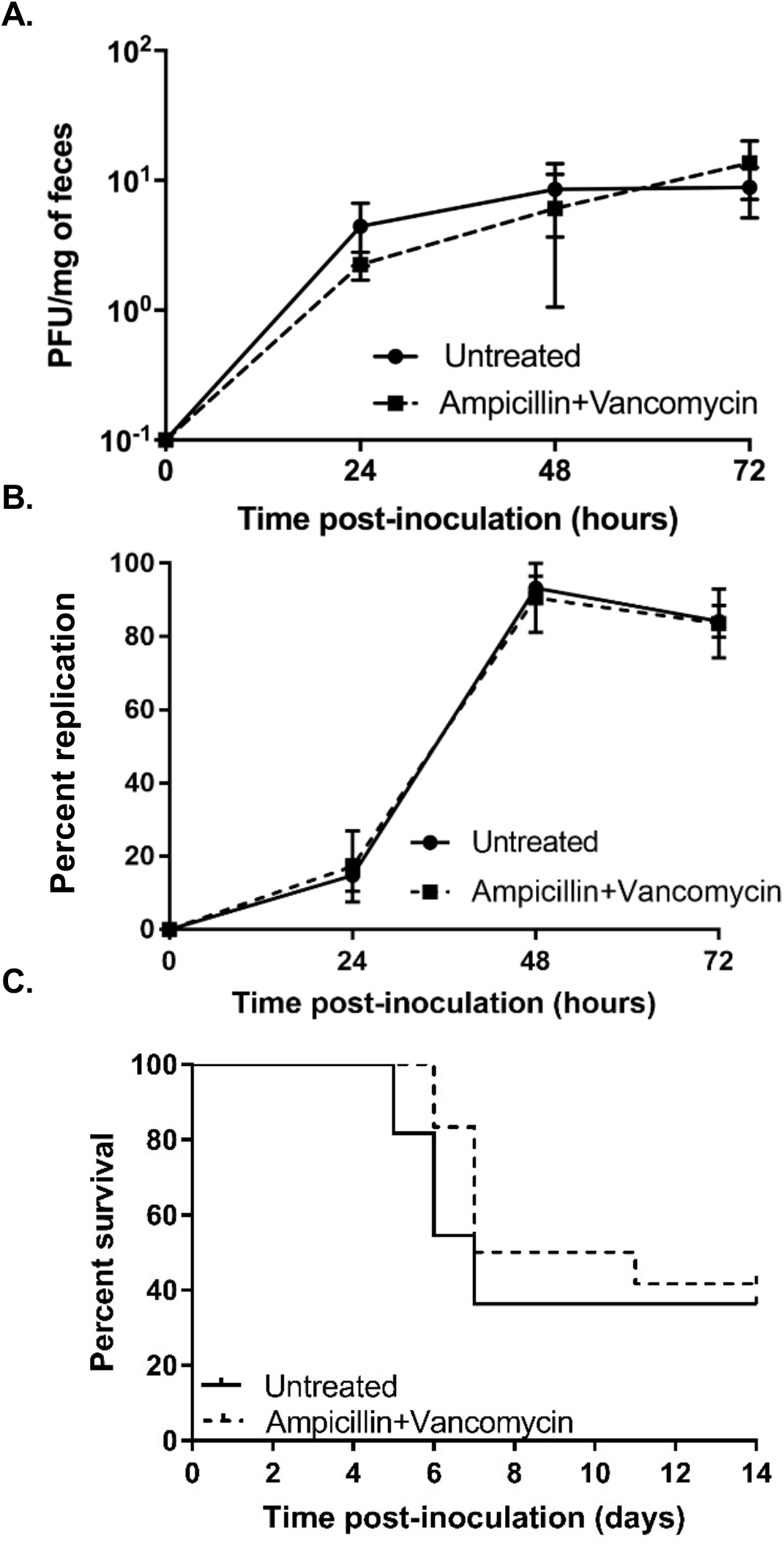
Treatment with ampicillin and vancomycin has no effect on poliovirus shedding and lethality. Male C57BL/6 PVR^+/+^ IFNAR^−/−^ mice were treated with or without various antibiotics for 5 days by oral gavage followed by *ad libitum* administration in drinking water for 9 days prior to oral inoculation with 2×10^7^ PFU of poliovirus. (A) Viral fecal titers at 24, 48, and 72 hours post inoculation were determined by plaque assay. (B) Viral replication efficiency was examined using light sensitive viruses. Mice were orally inoculated with neutral red-containing, light sensitive, poliovirus in the dark and feces were collected at 24, 48, and 72 hours post inoculation in the dark. Replication status was determined by dividing the number of PFU/ml of light-exposed samples (representing replicated virus) by the number of PFU/ml of dark-exposed samples (representing input/non-replicated virus plus replicated virus) and multiplying by 100%. Data are mean ± SEM. (C) Survival of untreated or ampicillin and vancomycin-treated mice following inoculation with 2×10^7^ PFU of poliovirus. Log-rank test. n=11-12 mice per group.

### Microbiota composition of mice treated with different antibiotic regimens

To examine the intestinal bacterial community of mice, we analyzed fecal samples using real-time quantitative PCR (qPCR) and 16S rRNA sequencing. As expected, qPCR experiments revealed that treatment with the five antibiotic cocktail or vancomycin and ampicillin significantly reduced prevalence of Eubacteria (general bacterial population, Fig. 4A), Bacteroidetes (a predominant phylum of Gram-negative bacteria, Fig. 4B), and Firmicutes (a predominant phylum of Gram-positive bacteria, Fig. 4C). Treatment with the five antibiotic cocktail reduced Eubacteria 16S rRNA gene copies over 80,000-fold and treatment with ampicillin and vancomycin reduced Eubacteria 16S rRNA gene copies by over 1,200-fold, and trends were similar for Bacteroidetes and Firmicutes (Fig. 4A-C). As expected, the five antibiotic cocktail depleted bacteria more than the ampicillin and vancomycin combination. To identify the microbial communities in each treatment group, we performed 16S rRNA sequencing. We found that the overall diversity of bacteria decreased in both mice treated with the five antibiotic cocktail and mice treated with ampicillin plus vancomycin (Fig. 4D). While microbial communities differed in mice treated with the five antibiotic cocktail vs. ampicillin and vancomycin (Fig. 4E), the relative prevalence of distinct members of the microbiota were more similar in the two antibiotic treatment groups than untreated controls (Fig. 4F). For example, in untreated mice, Bacteroidetes and Firmicutes were the most prevalent, while in the two different antibiotic treatment groups Tenericutes predominated. Overall, these data suggest that both antibiotic treatment regimens alter host microbiota, but that treatment with the cocktail of antibiotics depletes microbiota to a greater extent than treatment with ampicillin and vancomycin.

**Figure 4.**
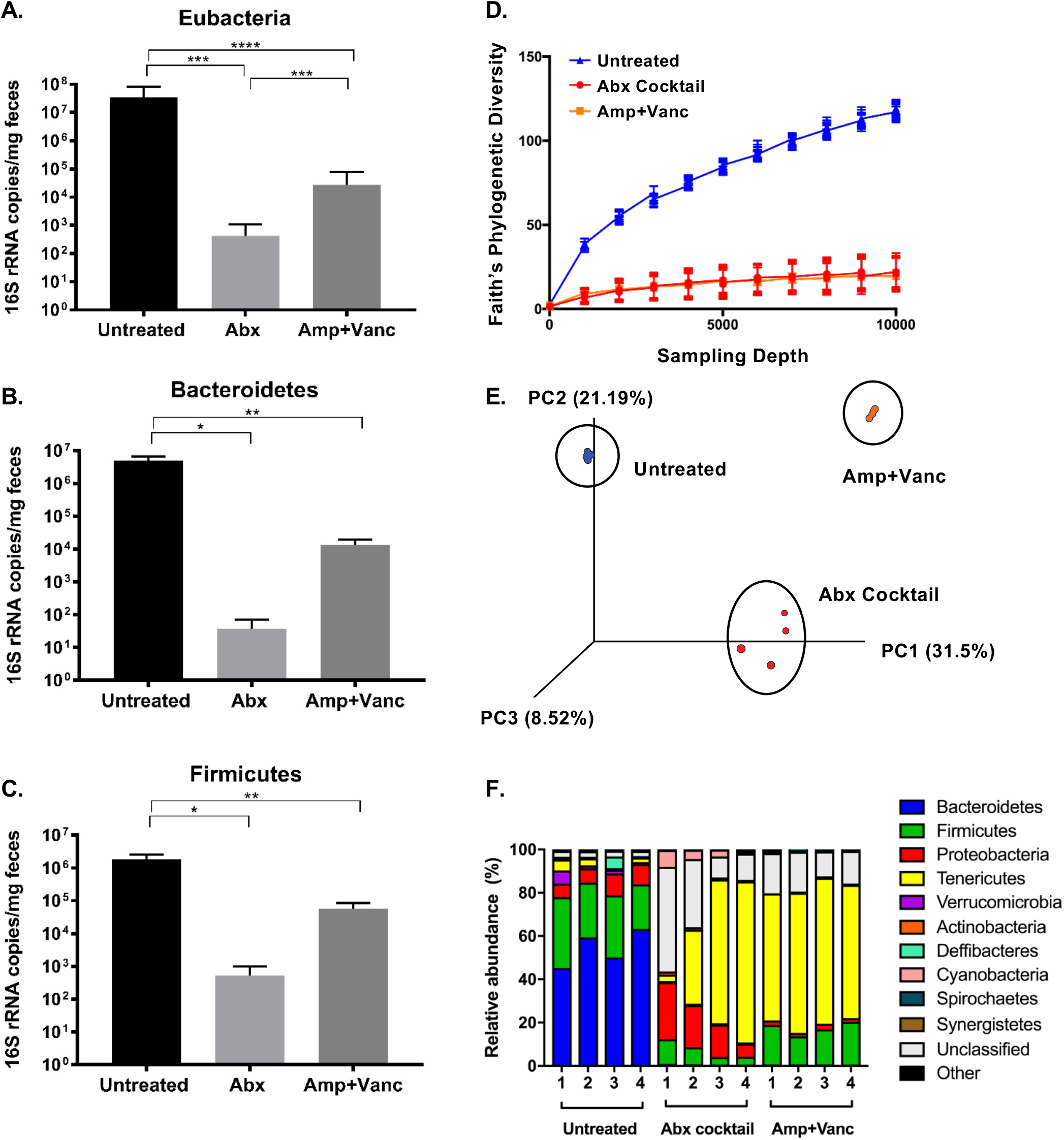
Profiling bacterial populations present in mice treated with different antibiotic regimens. Male C57BL/6 PVR^+/+^ IFNAR^−/−^ mice were treated with or without various antibiotics for 5 days by oral gavage followed by *ad libitum* administration in drinking water for 9 days. DNA was extracted from fecal pellets and bacteria were quantified by 16S rRNA qRT-PCR (A-C) and population diversity was determined by 16S rRNA gene sequencing (D-F). Group qRT-PCR (copies per mg of feces) performed on fecal genomic DNA collected from mice that were untreated, treated with a cocktail of five antibiotics, or treated with ampicillin and vancomycin: (A) Eubacteria (general bacterial population), (B) Bacteroidetes (a predominant phylum of Gram negative bacteria), and (C) Firmicutes (a predominant phylum of Gram positive bacteria). Data are mean ± SEM. *p<0.05, **p<0.001, ***p<0.0001, ****p<0.00001 Mann-Whitney Test. The V3-V4 region was amplified and sequenced using Illumina sequencing and analysis was performed on forward reads using Qiime: (D) Faith’s Phylogenetic Diversity (intrasample) using a rarefaction cutoff of 10000 reads/sample, (E) A principal component analysis plot was generated using Emporer (32), and (F) Bacterial taxonomy was assigned and visualized at level of class.

### CVB3, but not poliovirus, is sensitive to relatively minor perturbations of the microbiota

Given that treatment with just ampicillin and vancomycin was sufficient to inhibit CVB3 infection, we hypothesized that microbiota enhancement of CVB3 infection may require relatively high numbers of bacteria. To test this, we treated mice with milder antibiotic regimens. First, we treated mice with one-tenth of the normal dose (0.1X) of the five antibiotic cocktail and then orally infected them with either 5×10^7^ PFU of CVB3 or 2×10^7^ PFU of poliovirus. Feces were harvested at 72 hours post-inoculation and quantified via plaque assay. We found that treatment with the 0.1X five antibiotic cocktail was sufficient to significantly reduce CVB3 shedding, while having no effect on poliovirus shedding (Fig. 5A). To further examine the relative dependence of CVB3 and poliovirus on microbiota, we treated mice with a single dose of one antibiotic, streptomycin, the day prior to oral inoculation with 5×10^7^ PFU of CVB3 or 2×10^7^ PFU of poliovirus. Feces were harvested at 72 hours post-inoculation and viral titers were quantified via plaque assay. We found that mice treated with one dose of streptomycin had significantly reduced CVB3 shedding, while poliovirus shedding was unaffected (Fig. 5B). Furthermore, a single dose of streptomycin was sufficient to completely rescue mice from CVB3-associated lethality (Fig. 5C), while there was no effect on poliovirus-associated lethality (Fig. 5D). These data suggest that CVB3 is more sensitive to perturbations in the microbiota and that a single dose of one antibiotic can dramatically alter the outcome of infection. Overall, our data suggest that these two closely related enteric viruses have different microbiota requirements and mechanisms of microbiota utilization.

**Figure 5.**
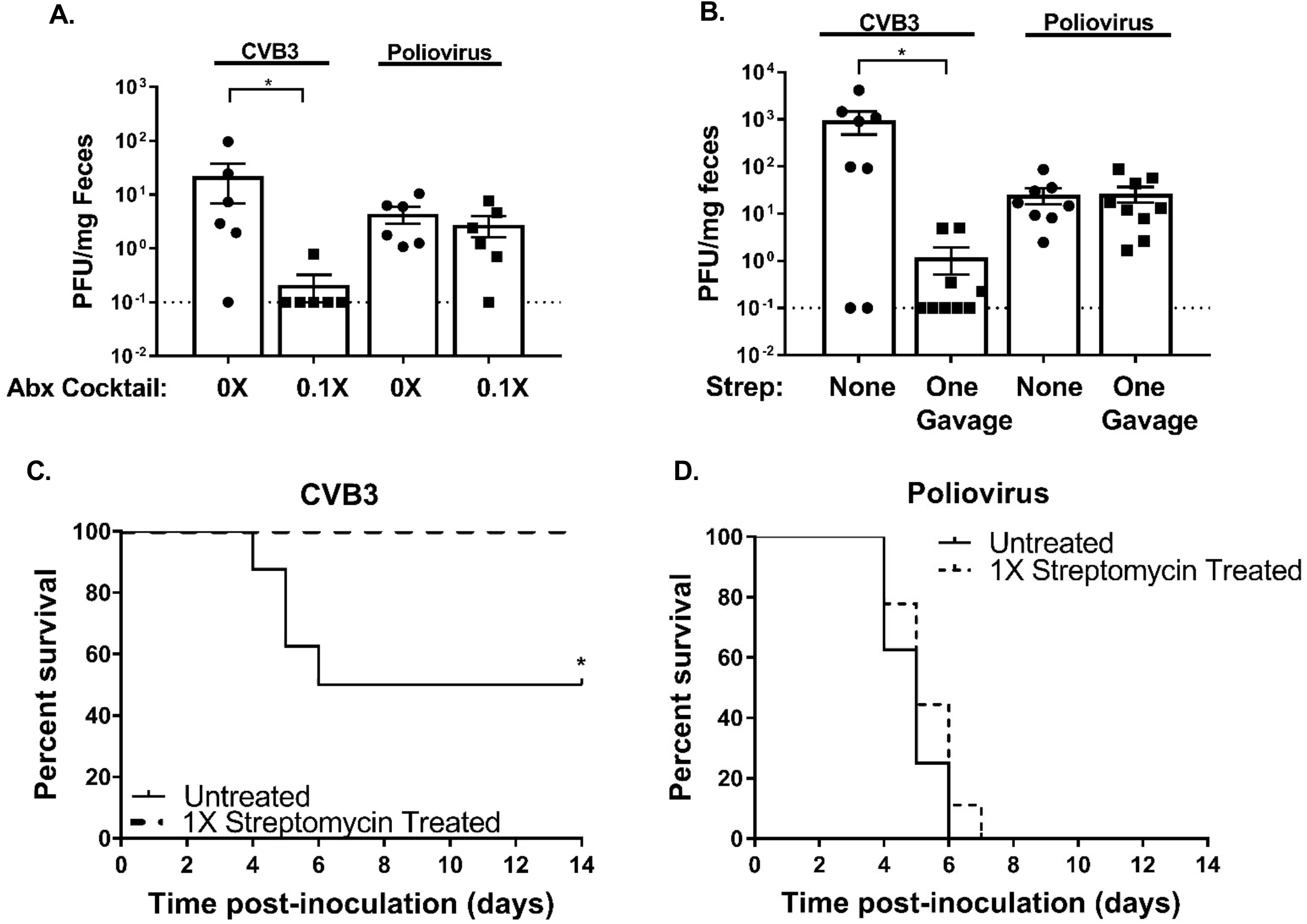
CVB3 replication is more sensitive to antibiotic treatment than poliovirus. (A) Male C57BL/6 PVR^+/+^ mice were treated with or without a combination of 5 antibiotics (ampicillin, neomycin, streptomycin, metronidazole, and vancomycin) at 0.1X concentration (one-tenth the standard concentration) for 5 days by oral gavage followed by 0.1X concentration *ad libitum* administration in drinking water for 9 days prior to oral inoculation with either 5×10^7^ PFU of CVB3 or 2×10^7^ of poliovirus. (A) Fecal titers at 72 hours post-inoculation were determined by plaque assay. (B-D) Male C57BL/6 PVR^+/+^ IFNAR^−/−^ mice were treated with or without a single gavage of streptomycin the day prior to oral inoculation with either 5×10^7^ PFU CVB3 or 2×10^7^ PFU poliovirus. (B) CVB3 and poliovirus fecal titers at 72 hours post inoculation were determined by plaque assay. Survival of untreated or streptomycin treated mice orally inoculated with CVB3 (C) or poliovirus (D). Data are mean ± SEM. *p<0.05 Mann-Whitney Test. *p<0.05, Log-rank test. n=6-9 mice per group.

## Discussion

Work from several labs has shown that enteric viruses from four families benefit from intestinal bacteria, but precise microbiota requirements for different viruses are unclear (6–9). Here we examined the role of bacteria on intestinal replication and pathogenesis of coxsackievirus, an enteric virus from the *Picornaviridae* family, using an oral inoculation mouse model. Not surprisingly, we found that CVB3 infection is sensitive to antibiotic-depletion of microbiota (Fig. 1). This suggests that, similar to poliovirus, bacteria promote CVB3 replication in the intestine.

Interestingly, we found that CVB3 is sensitive to relatively minor perturbations of the microbiota, whereas poliovirus is unaffected by these conditions and can replicate and cause disease. For example, CVB3 replication and pathogenesis was dramatically diminished by treatment with ampicillin and vancomycin (Fig. 2), treatment with a lower concentration of antibiotics (Fig. 5A), or even a single dose of a single antibiotic (Fig. 5B-D). Poliovirus replication and pathogenesis was unaffected by any of these treatment regimens. This suggests that even closely related enteric viruses may utilize different bacteria and/or mechanisms to enhance intestinal replication.

It is possible that CVB3 replication is dependent upon having a certain minimum number of bacteria present, having a particular bacterial strain or constellation of bacterial strains present, or a combination of these factors. We examined bacterial prevalence by 16S rRNA qPCR analysis and we examined bacterial diversity by 16S rRNA sequence analysis. We found that the overall abundance and diversity of the microbiota were altered during antibiotic treatment (Fig. 4). The relative diversity of bacterial phyla was similar between the whole cocktail of antibiotics and just ampicillin and vancomycin (Fig. 4D-F), whereas the relative bacterial abundance was more dramatically altered between these treatment groups (Fig. 4A-C). Thus, it is possible that bacterial prevalence may matter more than bacterial community composition for CVB3 replication, although future experiments will be needed to test this hypothesis.

While the precise mechanisms underlying the different microbiota requirements of CVB3 and poliovirus are unclear, it is possible that CVB3 is more reliant on microbiota for dampening host immune responses or enhancing the infectivity of viral particles. Treatment with ampicillin and vancomycin was sufficient to reduce CVB3 infection, and Baldridge et al. have shown that ampicillin and vancomycin are sufficient to regulate the IFN-λ pathway and reduce the persistence of another enteric virus, murine norovirus (7). It remains unclear if the IFN-λ pathway plays a role in bacterial enhancement of CVB3 intestinal replication. Additionally, antibiotics can regulate the host interferon response (22) and can have non-canonical effects on enteric viruses (23).

In conclusion, we found that similar to poliovirus and other recently identified enteric viruses, bacteria enhance CVB3 replication and viral-associated lethality. Interestingly, a single dose of a clinically relevant broad-spectrum antibiotic, streptomycin, was sufficient to reduce CVB3 replication and completely abolish CVB3-associated lethality, while having no effect on poliovirus. This result could have implications for how standard antibiotic treatment regimens for bacterial infections could also impact the course of enteric virus infections.

## Materials and Methods

### Cells and viruses

HeLa cells were propagated in Dulbecco’s modified Eagle’s medium (DMEM) supplemented with 10% calf serum. Poliovirus work was performed in BSL2+ areas in accordance with practices recommended by the World Health Organization. Poliovirus (serotype 1 virulent Mahoney) infections and plaque assays were performed using HeLa cells according to previously published methods (24). The CVB3-Nancy infectious clone was obtained from Marco Vignuzzi (Pasteur Institute, Paris, France). Stocks of CVB3 were prepared as previously described (20). CVB3 titers were determined by plaque assay with HeLa cells (20, 21).

To determine whether fecal viruses were input/inoculum virus or virus that had undergone replication, we used neutral-red-labeled, light-sensitive poliovirus as described previously (24). Light inactivation was performed by exposing neutral red-poliovirus stocks to a fluorescent light bulb for 10 minutes. The ratio of light-insensitive to light-sensitive PFU in the virus stock was 1 : 1.5×10^5^ PFU. Following oral inoculation, to determine the percentage of replicated virus in the intestine, samples were processed in the dark and a portion was light exposed. The percentage of replicated virus was calculated by dividing the light-exposed number of PFU/ml by the non-light-exposed number of PFU/ml and multiplying by 100.

### Mouse Experiments

All animals were handled according to the Guide for the Care of Laboratory Animals of the National Institutes of Health. All mouse studies were performed at UT Southwestern (Animal Welfare Assurance #A3472-01) using protocols approved by the local Institutional Animal Care and Use Committee in a manner designed to minimize pain, and any animals that exhibited severe disease were euthanized immediately with isoflurane. C57BL/6 PVR^+/+^ IFNAR^−/−^ mice were obtained from S. Koike (Tokyo, Japan) (25). Mice were administered either a combination of 5 antibiotics (ampicillin, neomycin, streptomycin, metronidazole, and vancomycin; 10mg of each antibiotic per day) or individual antibiotics (10mg per antibiotic per day) for 5 days by oral gavage followed by *ad libitum* administration in drinking water (ampicillin, neomycin, streptomycin, metronidazole: 1g/L; vancomycin: 500mg/L) for the duration of the experiment. Mice were treated with antibiotics for 9 days prior to peroral inoculation with poliovirus or CVB3. For oral inoculations, 8-12 week old male mice were perorally inoculated with either 5×10^7^ PFU of CVB3 or 2×10^7^ PFU of poliovirus. Male mice were used since CVB3 was shown to have a sex-dependent enhancement of replication and lethality (20). Disease was monitored until day 14 post-inoculation for survival experiments. Mice were euthanized upon severe disease onset. For viral shedding and replication experiments, feces were collected at indicated time points post-inoculation and processed as previously described (9). Tissues were aseptically removed from mice and homogenized in PBS using 0.9- to 2.0-mm stainless steel beads in a Bullet Blender (Next Advance) and freeze-thawed 3 times in liquid nitrogen to release intracellular virus. Cellular debris was removed by centrifugation at 13,000 RPM for 3 minutes. The amount of virus in supernatant was determined by plaque assay in HeLa cells and titers were normalized to the weight of each tissue.

### 16S qRT-PCR and sequencing

DNA was extracted from fecal pellets using PowerFecal DNA (Qiagen). The V3-V4 region of the 16S rRNA gene was amplified using primers, Forward primer: 5’ TCGTCGGCAGCGTCAGATGTGTATAAGAGACAG-[CCTACGGGNGGCWGCAG] Reverse primer: 5’ GTCTCGTGGGCTCGGAGATGTGTATAAGAGACAG-[GACTACHVGGGTATCTAATCC] and sequenced on Illumina MiSeq v3 using PE 300 Cycle. Analysis was performed on forward reads using Qiime 1.9.1 (26). Primer sequences were trimmed from each read and all reads were quality filtered (Phred score > 25) using the multiple_split_libraries_fastq.py command. Operational taxonomic units (OTUs) were assigned using open-reference picking (SILVA 128 release) (27, 28) using default parameters, except for 0.1% subsampling, and for filter_alignment.py: gap filter threshold, 0.8; lane mask filtering, true; entropy threshold, 0.10. This yielded a minimum of 134043 and median of 238345 sequence counts per sample. Bacterial taxonomy was assigned (29, 30) and visualized at level of class. Intra-sample and between sample diversity was calculated using core_diversity.py workflow (31). Faith’s Phylogenetic Diversity (intra-sample) used a rarefaction cutoff of 10000 reads/sample. PCoA plot was visualized using Emporer (32).

#### Statistical Analysis

The difference between groups were examined by the unpaired two-tailed Student *t* test. Error bars in figures represent the standard errors of the means. A *P* value of <0.05 was considered significant. All analyses of data were performed using GraphPad Prism version 7.00 for Windows, GraphPad Software, La Jolla, CA.

## Acknowledgements

This work is funded by a Burroughs Wellcome Fund Investigator in the Pathogenesis of Infectious Diseases award (J.K.P.), R01 AI74668 (J.K.P.), and K01 DK110216 (C.M.R.). J.K.P. is a Howard Hughes Medical Institute Faculty Scholar. MWA was supported in part by NIH NIAID grant T32 AI007520. BM was supported by NIH NIAID grant T32 AI007520 and F32 AI138392.

